# Four chemoreceptors govern bidirectional pH taxis in *Bacillus subtilis*

**DOI:** 10.1101/715946

**Authors:** Payman Tohidifar, Matthew J. Plutz, George W. Ordal, Christopher V. Rao

**Affiliations:** Department of Chemical and Biomolecular Engineering, University of Illinois at Urbana-Champaign, Urbana, Illinois, USA; Department of Biochemistry, University of Illinois at Urbana-Champaign, Urbana, Illinois, USA

## Abstract

We investigated pH taxis in *Bacillus subtilis*. This bacterium was found to perform bidirectional taxis in response to external pH gradients, enabling it to preferentially migrate to neutral environments. We next investigated the chemoreceptors involved in sensing pH gradients. We found that four chemoreceptors are involved in sensing pH: McpA and TlpA for sensing acidic environments and McpB and TlpB for alkaline ones. In addition, TlpA was found to also weakly sense alkaline environments. By analyzing chimeras between McpA and TlpB, the principal acid and base-sensing chemoreceptors, we identified four critical amino-acid residues – Thr^199^, Gln^200^, His^273^, and Glu^274^ on McpA and Lys^199^, Glu^200^, Gln^273^, and Asp^274^ on TlpB – involved in sensing pH. Swapping these four residues between McpA and TlpB converted the former into a base receptor and the latter into an acid receptor. Based on the results, we propose that disruption of hydrogen bonding between the adjacent residues upon pH changes induces signaling. Collectively, our results further our understanding of chemotaxis in *B. subtilis* and provide a new model for pH sensing in bacteria.

**IMPORTANCE:** Many bacteria can sense the pH in their environment and then use this information to direct their movement towards more favorable locations. In this study, we investigated the pH sensing mechanism in *Bacillus subtilis*. This bacterium preferentially migrates to neutral environments. It employs four chemoreceptors to sense pH. Two are involved in sensing acidic environments and two are involved in sensing alkaline ones. To identify the mechanism for pH sensing, we constructed receptor chimeras of acid and base sensing chemoreceptors. By analyzing the response of these chimeric receptors, we were able to identify four critical amino-acid residues involved in pH sensing and propose a model for the pH sensing mechanism in *B. subtilis*.

## INTRODUCTION

*Bacillus subtilis* performs chemotaxis to a wide range of attractants and repellents (1–3). As a brief background, *B. subtilis* employs ten chemoreceptors to sense these compounds (4). The governing chemotaxis pathway functions differently than the better understood chemotaxis pathway in *Escherichia coli*. The core signaling pathways consists of a membrane-associated complex involving the chemoreceptors, CheA histidine kinase, and CheW and CheV scaffold proteins (5) that preferentially form clusters at the cell poles (6). The chemoreceptors sense attractants and repellents either by binding them directly or through binding proteins associated with their uptake (7). Attractants are known to increase the rate of CheA phosphorylation (8). The phosphate group is then transferred to CheY, which in the phosphorylated form binds to the cytoplasmic face of the flagellar motors and induces a motile response (9).

To sense chemical gradients, *B. subtilis* employs three adaptation systems (10–12). The primary system involves receptor methylation. Two enzymes, the CheR methyltransferase and the CheB methylesterase, add and remove methyl groups on conserved glutamate residues located on the cytoplasmic domain of the chemoreceptors (13, 14). Depending on the specific glutamate residue, these modifications can either increase or decrease the ability of the chemoreceptors to activate the CheA kinase (5, 15). In addition, two other adaptation systems are involved in sensing gradients. One involves the scaffold protein CheV, which contains a C-terminal response regulator domain that is phosphorylated by CheA (16). Depending on the methylation state of the chemoreceptors, phosphorylated CheV can either increase or decrease chemoreceptor activity (5). The other system involves three proteins: CheD, a chemoreceptor deamidase (17, 18); CheC, a phosphatase for phosphorylated CheY (19); and CheY. In addition to being a deamidase, CheD binds the chemoreceptors and increases their ability to activate CheA (5, 20). CheC can also bind CheD and prevent it from binding the chemoreceptors. Phosphorylated CheY increases the affinity between CheC and CheD, thus providing a feedback mechanism for controlling CheA activity in response to phosphorylated CheY (21, 22).

Our understanding of *B. subtilis* chemotaxis is principally limited to amino-acid chemotaxis. Aside from amino acids (7, 23), oxygen (24), and sugars transported by the phosphoenolpyruvate-dependent phosphotransferase system (25), little is known about the sensing mechanisms involved in *B. subtilis* chemotaxis. A number of reports have shown that diverse bacteria migrate in response to pH gradients. This process is best understood in *Escherichia coli, Salmonella enterica* and *Helicobacter pylori* (26–32). In the case of *E. coli* and *S. enterica*, these bacteria preferentially migrate to neutral (pH 7.5) environments (33). The response is bidirectional, as the bacteria will migrate down pH gradients when the ambient pH is too high or migrate up pH gradients with the ambient pH is too low. The underlying mechanism involves the competitive response between two chemoreceptors, one that induces cells to migrate down pH gradients and the other that induces them to migrate up pH gradients. These two chemoreceptors respond to both internal and external pH. While the sensing mechanism is still not well understood, external pH is believed to be sensed by the extracellular domain of the chemoreceptors (27) and internal pH by the linker region between the transmembrane helices and cytoplasmic domain of the chemoreceptors (30). Swapping the entire linker region or specific amino-acid residues within this linker region inverts the response of these two chemoreceptors to changes in internal pH (30).

In this work, we investigated chemotaxis to external pH gradients in *B. subtilis*. Similar to *E. coli* and *S. enterica, B. subtilis* exhibits bidirectional chemotaxis to pH gradients. To sense these pH gradients, *B. subtilis* employs four chemoreceptors, two for sensing acids and two for sensing base. By analyzing chimeras between acid and base-sensing chemoreceptors, we identified four critical amino-acid residues involved in sensing external pH. Swapping these four residues changed a base-sensing chemoreceptor into an acid-sensing one, and vice versa. Based on these data, we propose a model for pH sensing in *B. subtilis*.

## RESULTS AND DISCUSSION

### *B. subtilis* exhibits bidirectional taxis to external pH gradients

To determine whether *B. subtilis* performs chemotaxis in response to external pH gradients, we employed the capillary assay (26). Briefly, cells suspended in buffer at different pH’s (6.0-8.5) were incubated with capillaries filled with buffer at pH 7.0 and then the number of cells that entered the capillaries after 1 h were counted. The resulting data show that *B. subtilis* exhibits bidirectional taxis to pH gradients in manner similar to what is observed in *E. coli* (**Figure 1A**). In particular, we found that *B. subtilis* preferentially migrates to neutral (pH 7) environments when the cells were initially suspended in either acidic or alkaline buffer (pH 6-8). Outside of this pH range, however, the cells were less motile (data not shown) and, consequently, taxis was reduced.

**Figure 1.**
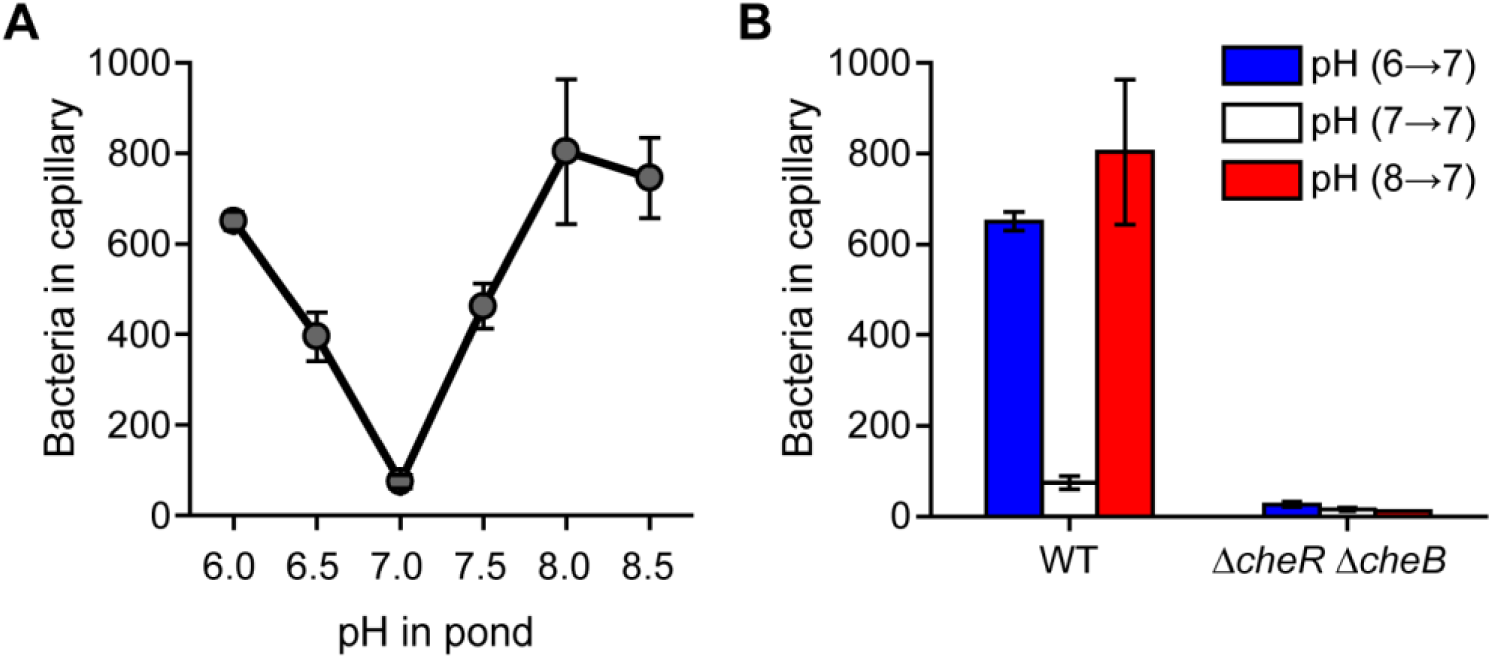
*B. subtilis* exhibits bidirectional chemotaxis to pH gradients. **(A)** Response to increasing and decreasing pH gradients. **(B)** Methylation is necessary for pH taxis.

### The methylation system is required for pH taxis

*B. subtilis* employs three adaptions systems for sensing chemical gradients (11). Of the three adaptation systems, the methylation system is the dominant one for sensing amino-acid gradients. To determine whether the methylation system is also required for pH taxis, we tested whether a Δ*cheR*Δ*cheB* mutant was capable of pH taxis (**Figure 1B**). This mutant lacks the requisite methyltransferase (CheR) and methylesterase (CheB). It was unable to perform pH taxis, indicating that the methylation system is necessary for sensing pH gradients.

### Four chemoreceptors are involved in sensing pH gradients

*B. subtilis* possesses ten chemoreceptors (4). To determine which chemoreceptors are involved in pH taxis, we tested mutants expressing just one chemoreceptor (**Figure 2A**). Of the single chemoreceptor mutants, only the one expressing McpA as its sole chemoreceptor was capable of acid sensing. In addition, this mutant exhibited a weak repellent response to base, with fewer cells migrating into the capillary when the buffer was at pH 6 as compared to pH 7 (33.7 ± 10.5 versus 54.7 ± 7.1 cells). This repellent response is consistent with McpA being an acid sensor. Two single chemoreceptor mutants were found to exhibit base sensing: the strains expressing McpB or TlpB as their sole chemoreceptors. In particular, these strains exhibited taxis to increases in pH. They also exhibited a weak repellent response to acid (McpB: 37.3 ± 8.6 versus 51.0 ± 15.1 cells; TlpB: 44.3 ± 2.5 versus 68.7 ± 15.8 cells). No significant responses to any changes in pH were observed for the other mutants. The existence of chemoreceptors sensing both acids and bases would explain how *B. subtilis* is capable of preferentially swimming to neutral conditions, namely that it reflects competition between the acid and base responses.

**Figure 2.**
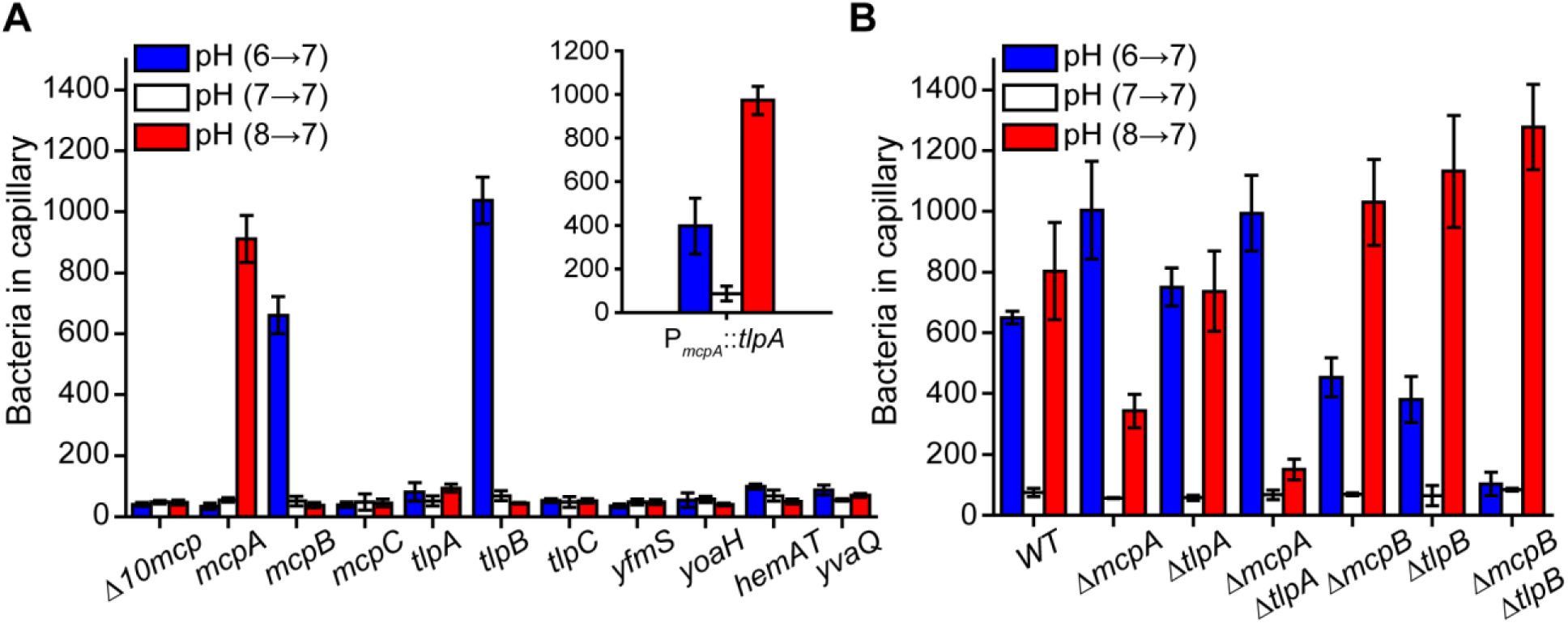
McpA and TlpA are the principle chemoreceptors involved in the acid response and McpB and TlpB are the sole chemoreceptors involved in the base response. **(A)** Response of strains expressing just one chemoreceptor to pH gradients. **Inset**: response of strain over-expressing *tlpA* as its sole chemoreceptor. **(B)** Response of mutants lacking key receptors to pH gradients. Error bars denote the standard deviation of three biological replicates performed on three separate days.

The genes encoding McpA, McpB, TlpA, and TlpB reside in a four-gene cluster (34). Since these four chemoreceptors exhibit high (57-65%) amino-acid sequence identify, we hypothesized that TlpA may also be involved even though a strain expressing it as the sole chemoreceptor failed to exhibit a response to changes in pH. The reason may be that TlpA is weakly expressed: the wild type expresses 2,000 copies of this chemoreceptor as compared to 16,000 copies for McpA (35). Therefore, we tested whether expressing TlpA from a stronger promoter would enable pH sensing. When *tlpA* was expressed as the sole chemoreceptor using the *mcpA* promoter, we observed both an acid and base response (**Figure 2A**, inset). The acid response, however, was stronger than the base response. These results imply that TlpA alone, when over expressed, can direct *B. subtilis* to neutral environments. This begs the question as to why multiple chemoreceptors are employed for pH taxis when potentially one would suffice. Unfortunately, we cannot answer this question at this time.

We next tested the effect of deleting these four chemoreceptors, both individual and in combination, on pH sensing (**Figure 2B**). When *mcpA* was deleted in the wild type (Δ*mcpA*), we observed a reduction in acid sensing, consistent with this chemoreceptor being involved in chemotaxis towards lower pH’s. We also observed an increase in the base response, perhaps reflecting the competition between the acid and base responses. When *tlpA* was deleted in the wild type (Δ*tlpA*), we observed no significant change in pH sensing. However, when both of the chemoreceptors were deleted in the wild type (Δ*mcpA*Δ*tlpA*), the acid response was almost completely eliminated and the base response increased. These results suggest that McpA and TlpA are the primary chemoreceptors involved in sensing decreases in pH. Additional chemoreceptors may be involved; however, their contribution appears minor.

When either McpB or TlpB were deleted in the wild type (Δ*mcpB* or Δ*tlpB*), the base response was reduced. In addition, the acid response increased. When both chemoreceptors were deleted (Δ*mcpB*Δ*tlpB*), the base response was completely eliminated and the acid response further increased. These results suggest that McpB and TlpB are the sole chemoreceptors involved in sensing increases in pH.

### Identification of the regions involved in sensing pH gradients

All four chemoreceptors involved in pH taxis employ the same double Cache domain (dCache_1) for their extracellular sensing domain (36). This would suggest that specific amino-acid residues are involved in pH. As a first step towards identifying these residues, we constructed chimeras between McpA, involved in acid sensing, and TlpB, involved in base sensing. We focused on these two chemoreceptors because they are primary ones involved in pH taxis. One challenge with constructing these chimeras is that they may not be functional. Indeed, many were not. Unfortunately, little was known prior regarding any additional ligands for these chemoreceptors. As *B. subtilis* is known to perform chemotaxis towards amino acids, we tested whether these chemoreceptors were involved in sensing casamino acids. TlpB alone was able to support chemotaxis to casamino acid; however, McpA alone did not. While we were not able to identify any attractants for McpA, we did find that McpA governs the repellent response to indole.

We first constructed a series of chimeras where we fused the N-terminal region of TlpB with the C-terminal region of McpA: *tlpB*_362_*mcpA, tlpB*_*284*_*mcpA, tlpB*_*260*_*mcpA, tlpB*_*238*_*mcpA* and *tlpB*_*180*_*mcpA* (**Figure 3**). We then tested the ability of strains expressing these hybrids as their sole chemoreceptor to sense acid and base gradients using the capillary assay. In addition, we employed casamino acids as a control. Strains expressing *tlpB*_362_*mcpA* or *tlpB*_*284*_*mcpA* behaved the same as wild-type *tlpB* (**Figure 4**). These results indicate that pH is not sensed by the cytoplasmic domain of the chemoreceptor or by the HAMP domain. The strain expressing *tlpB*_*260*_*mcpA* was able to sense both increases and decreases in pH. As the base response was similar to the strain expressing wild-type *tlpB*, these results would suggest that the region 260-284 from McpA is involved in acid sensing. The strain expressing *tlpB*_*238*_*mcpA* was able to sense both increases and decreases in pH, albeit at reduced levels. However, the strains expressing *tlpB*_*180*_*mcpA* were unable to sense base gradients and only responded to acid gradients, indicating that the region spanning the residues 180-284 is directly involved in pH sensing. In addition, strains expressing *tlpB*_*180*_*mcpA* no longer responded to casamino acids. Likely, this is due to disruption of the sensing domain. All other N-terminal TlpB chimeras were able to sense casamino-acid gradients.

**Figure 3.**
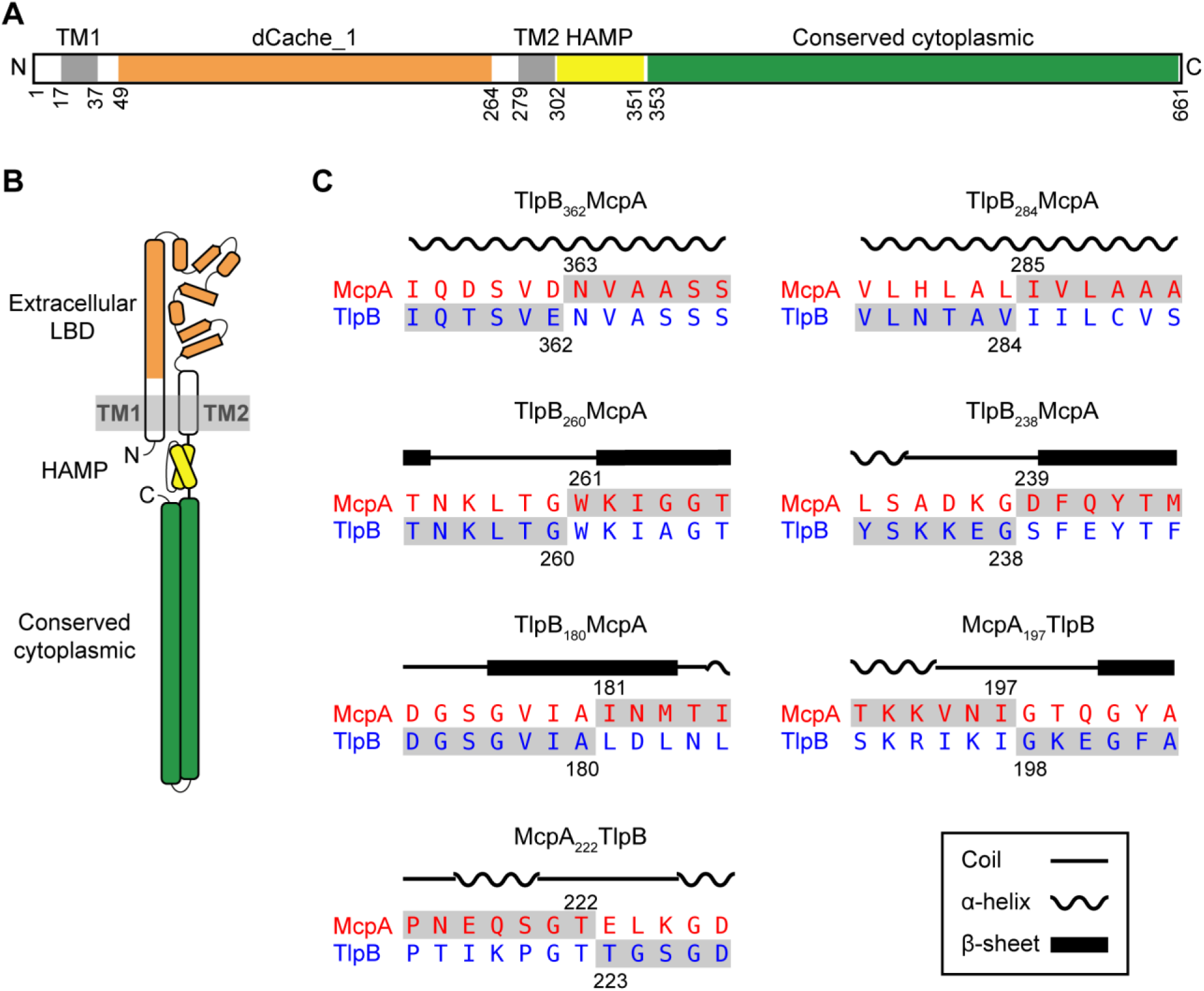
Construction of chimeric receptors to determine regions involved in pH sensing. **(A)** Domain structure of McpA. Extracellular ligand-binding, dCache_1 domain (orange), transmembrane transmembrane, TM1 and TM2 (gray), HAMP domain (yellow), and cytoplasmic domain (green). **(B)** Cartoon structure of McpA. **(C)** Amino-acid sequence alignment of McpA and TlpB around chimera junction points. The numbers designate the fusion points between two chemoreceptors, and the local sequences of the final chimeric chemoreceptors are highlighted in gray.

**Figure 4.**
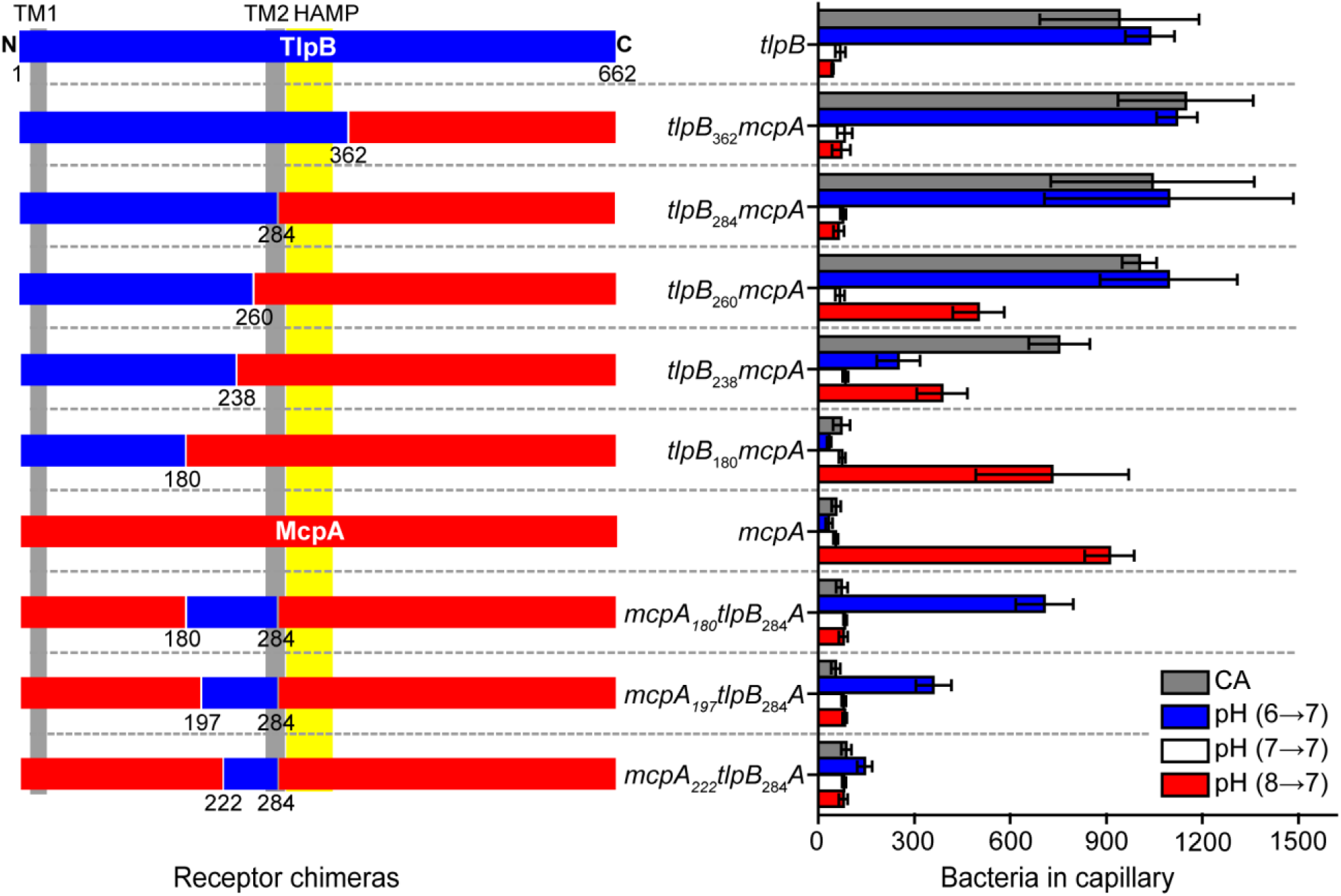
Response of strains expressing receptor chimeras involving different fragments of McpA (red) and TlpB (blue) to pH and casamino-acid (CA) gradients. Error bars denote the standard deviation of three biological replicates performed on three separate days.

To further characterize the region spanning residues 180-284, we replaced this region in McpA with the corresponding region from TlpB (*mcpA*_*180*_*tlpB*_*284*_*A*). Strains expressing this chimera as their sole chemoreceptor only responded to increases in pH (**Figure 4**). We then created two more chimeric strains, *mcpA*_*197*_*tlpB*_*284*_*A* and *mcpA*_*222*_*tlpB*_*284*_*A*, to narrow down the regions responsible for pH sensing. The mutant expressing *mcpA*_*197*_*tlpB*_*284*_*A* also responded only to increases in pH. However, the mutant expressing *mcpA*_*222*_*tlpB*_*284*_*A* did not exhibit a significant response to pH (**Figure 4**). These results indicated that the sub-region spanning the residues 197-222 is also involved in base sensing.

All of the strains expressing the McpA chimeras exhibited a repellent response to indole. However, the response of these strains was significantly reduced as compared to the strain expressing wild-type *mcpA* (average net accumulation within capillary: 219.7 ± 90.5 (*mcpA*_*180*_*tlpB*_*284*_*A*), 232.0 ± 183.1 (*mcpA*_*197*_*tlpB*_*284*_*A*), and 248.6 ± 109.5 (*mcpA*_*222*_*tlpB*_*284*_*A*) versus 1,286.1 ± 467.6 cells (wild type)). This would suggest that the McpA chimeras are not fully functional, which may explain their reduced response to increases in pH as compared to TlpB.

### Identification of residues involved in pH sensing

To identify the specific amino-acid residues involved in pH sensing, we aligned the amino-acid sequences spanning residues 195-284 on the four chemoreceptors (**Figure 5A**). Charged amino acids are more likely to be involved in pH sensing as their side-chains can be protonated and deprotonated. In addition, the polar amino acids may play an indirect role in pH sensing by impacting the local amino-acid p*K*_*a*_ values and/or forming hydrogen bonds with side-chains of ionizable amino-acid residues. We first focused on the charged amino acids that carried opposite signs on McpA and TlpB. Within the pH sensing sub-region spanning residues 260-284, the potential candidate residues were the Gln^273^-Asp^274^ pair for TlpB and the corresponding His^273^-Glu^274^ pair for McpA. Within the other pH sensing sub-region spanning residues 197-222, the potential candidate residues were the Ile^218^- Lys^219^ pair for TlpB and the corresponding Glu^218^-Gln^219^ pair for McpA.

**Figure 5.**
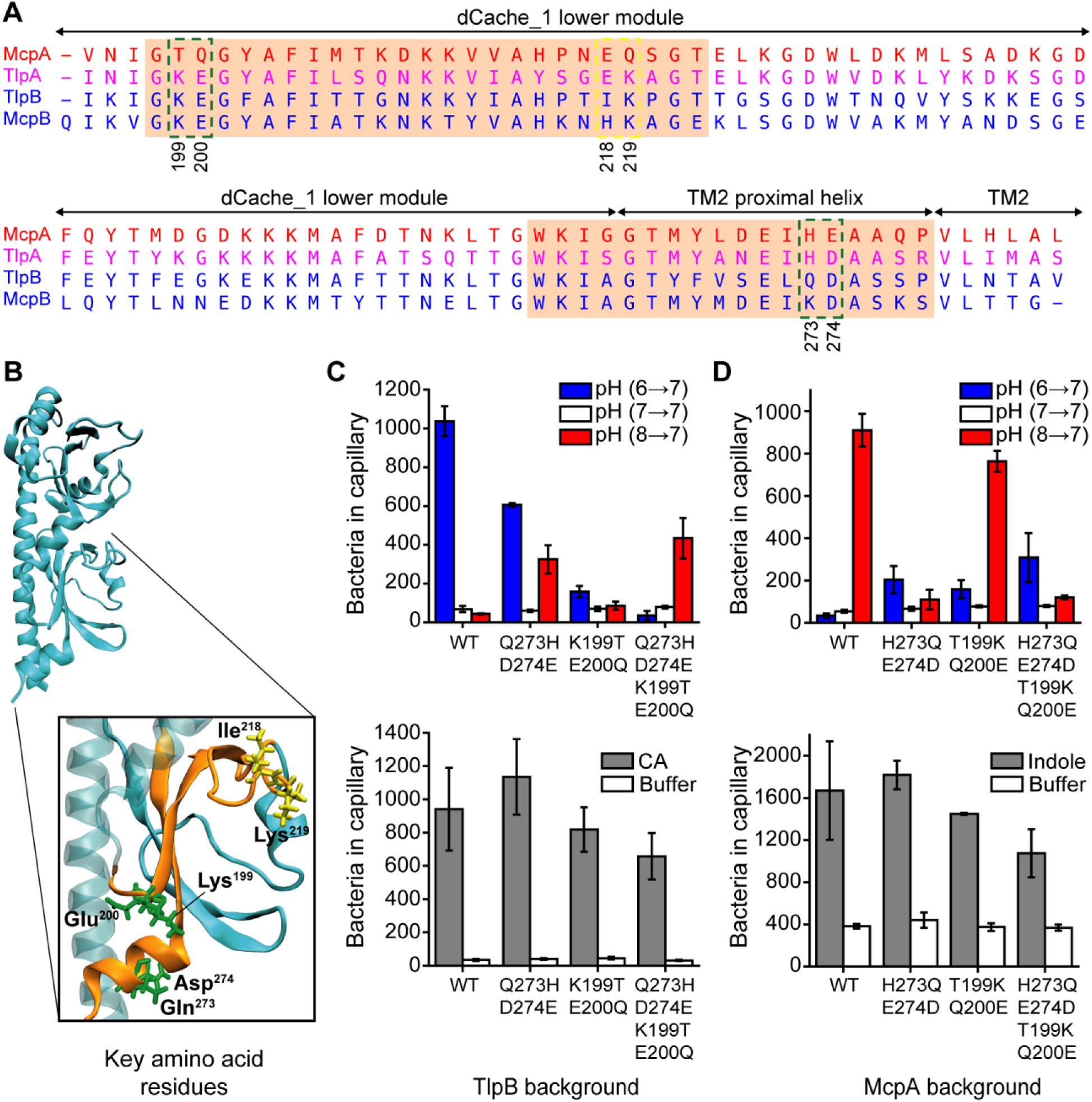
Identification of critical residues involved in sensing pH. **(A)** Amino-acid sequence alignment of pH sensing regions spanning residues (195-284) for the four pH chemoreceptors reveals candidate residues for mutational analysis. Candidate residues are shown within the green and yellow dashed boxes. Amino-acid sequence for TlpA are shown in purple because TlpA is sensitive to both acid and base. pH sensing sub-regions identified from chimeric receptor analysis are highlighted in the orange boxes. **(B)** Predicted structure of the TlpB ligand binding domain (LBD). Two pH sensing sub-regions are shown in orange consistent with panel A. The candidate amino-acid residues on the TlpB extracellular LBD are shown in green and yellow. (**C**) Response of strains expressing *tlpB* mutants to pH and casamino-acid (CA) gradients. (**D**) Response of strains expressing *mcpA* mutants to pH and negative indole gradients. Error bars denote the standard deviation of three biological replicates performed on three separate days.

The pH sensing sub-regions are separated by 38 amino-acid residues. However, when the predicted tertiary structure of TlpB sensing domain was visualized, we found the amino-acid pair Lys^199^-Glu^200^ was in close proximity to the Gln^273^-Asp^274^ pair (**Figure 5B**). Similarly, we observed the mutual Thr^199^-Gln^200^ pair was in close proximity to the His^273^-Glu^274^ pair on McpA (not shown in Figure). Therefore, Lys^199^-Glu^200^ on TlpB and Thr^199^-Gln^200^ on McpA were also potential candidate residues for mutational analysis as local amino-acid interactions could affect pH sensing.

We examined the role of these candidate pairs by swapping them with their counterparts on the opposite receptor, both individually and in combination. We first replaced Gln^273^-Asp^274^ pair on TlpB with the counterpart His^273^-Glu^274^ pair from McpA. Cells expressing *tlpB-Q273H/D274E* as their sole chemoreceptor exhibited a reduced base response and an increased acid response (**Figure 5C**). When we performed the reciprocal experiment, replacing the His^273^-Glu^274^ pair on McpA with the Gln^273^-Asp^274^ pair from TlpB, we observed a large decrease in the acid response and small increase in the base response (**Figure 5D**). These results demonstrate that Gln^273^-Asp^274^ on TlpB and His^273^-Glu^274^ on McpA are involved in pH sensing.

We next tested the effect of replacing the Lys^199^-Glu^200^ pair on TlpB with the Thr^199^- Gln^200^ pair from McpA. This mutant (*tlpB-K199T/E200Q*) exhibited a weak response to both acids and bases (**Figure 5C**). However, the reciprocal mutation (*mcpA- T199K/Q200E*), where we replaced the Thr^199^-Gln^200^ pair on McpA with the Lys^199^- Glu^200^ pair from TlpB, led to a small reduction in the acid response and a small increase in the base response (**Figure 5D**). When the four residues on TlpB were swapped with their counterpart residues for McpA, the base response was eliminated and the resulting strain only exhibited an acid response, albeit at reduced levels (**Figure 5C**). Similarly, when the four residues on McpA were swapped with their counterpart residues from TlpB, the acid response was significantly reduced and the resulting strain exhibited a base response, again at reduced levels. Collectively, these results imply that these four amino-acid residues are sufficient to define the polarity of pH sensing for both McpA and TlpB. Therefore, we did not examine Ile^218^-Lys^219^ pair on TlpB or Glu^218^-Gln^219^ on McpA as these residues are located far away from the identified key residues (**Figure 5B**) and likely are not involved in pH sensing.

We note that these mutant chemoreceptors with swapped polarity exhibited a reduced response to pH. Likely, the mutations are disrupting overall chemoreceptor function. When we tested the chemoreceptors against casamino acids and indole, they exhibited a weaker response than the wild type.

### Model for pH sensing

The experiments described above identified four critical amino-acid residues involved in pH sensing. The predicted structures for the extracellular domains of TlpB and McpA reveal that these four amino-acid residues are in close proximity of one another, suggesting that direct interactions between them may govern the sensing mechanism (see **Fig. 5B**). Side-chains of ionizable amino-acid residues can accept or donate protons as the local pH varies. It is generally thought that changes in protonation state of ionizable amino-acid residues can lead to conformational changes in the protein and possibly alter its activity. These conformational changes are typically induced by the formation or disruption of hydrogen bonds between two or more proximal residues. The protonation state of ionizable amino-acid residues largely relies on their local p*K*_*a*_ values. The local p*K*_*a*_ of an amino-acid residue in the folded protein depends on its interactions with neighbor residues. As the result of such interactions – including charge-dipole, charge-charge, and ion-pairs – p*K*_*a*_ values can be significantly different from the intrinsic p*K*_*a*_ (p*K*_*a*-int_) values measured in blocked pentapeptides (37). For example, the p*K*_*a*_ of a lysine residue can be as low as 5.7 and the p*K*_*a*_ of an aspartate residue can be as high as 9.2 in folded proteins (37). In the case of TlpB, increase in pH can directly affect three ionizable residues: Asp^274^, Lys^199^, and Glu^200^. The point mutation *tlpB-D274N* did not affect the responses to pH (1,334.7 ± 352.4 versus 1,037.2 ± 76.9 cell for wild-type *tlpB*), suggesting that the protonation state of Asp^274^ remains intact or does not affect interactions with other residues. However, the double mutant K199T/E200Q significantly reduced the response to increase in pH (**Figure 5C**). This result implies increases in pH affects the protonation state of Lys^199^ or/and Glu^200^. Based on these results, we offer one potential model to explain the response of TlpB to increases in pH (**Figure 6**). At lower pH (e.g. pH=6), Lys^199^ and/or Glu^200^ are protonated and form stable hydrogen bonds with Gln^273^ or/and Asp^274^. As pH increases, Lys^199^ and/or Glu^200^ likely become deprotonated, and the loss of the hydrogen bonds destabilizes the local structure of lower region of the sensing domain and the proximal transmembrane helix (TM2, **Fig. 3**). This structural transition is then propagated through TM2 to the HAMP domain and subsequently to distal cytoplasmic signaling, thus inducing the autophosphorylation of CheA histidine kinase (**Figure 6A**).

**Figure 6.**
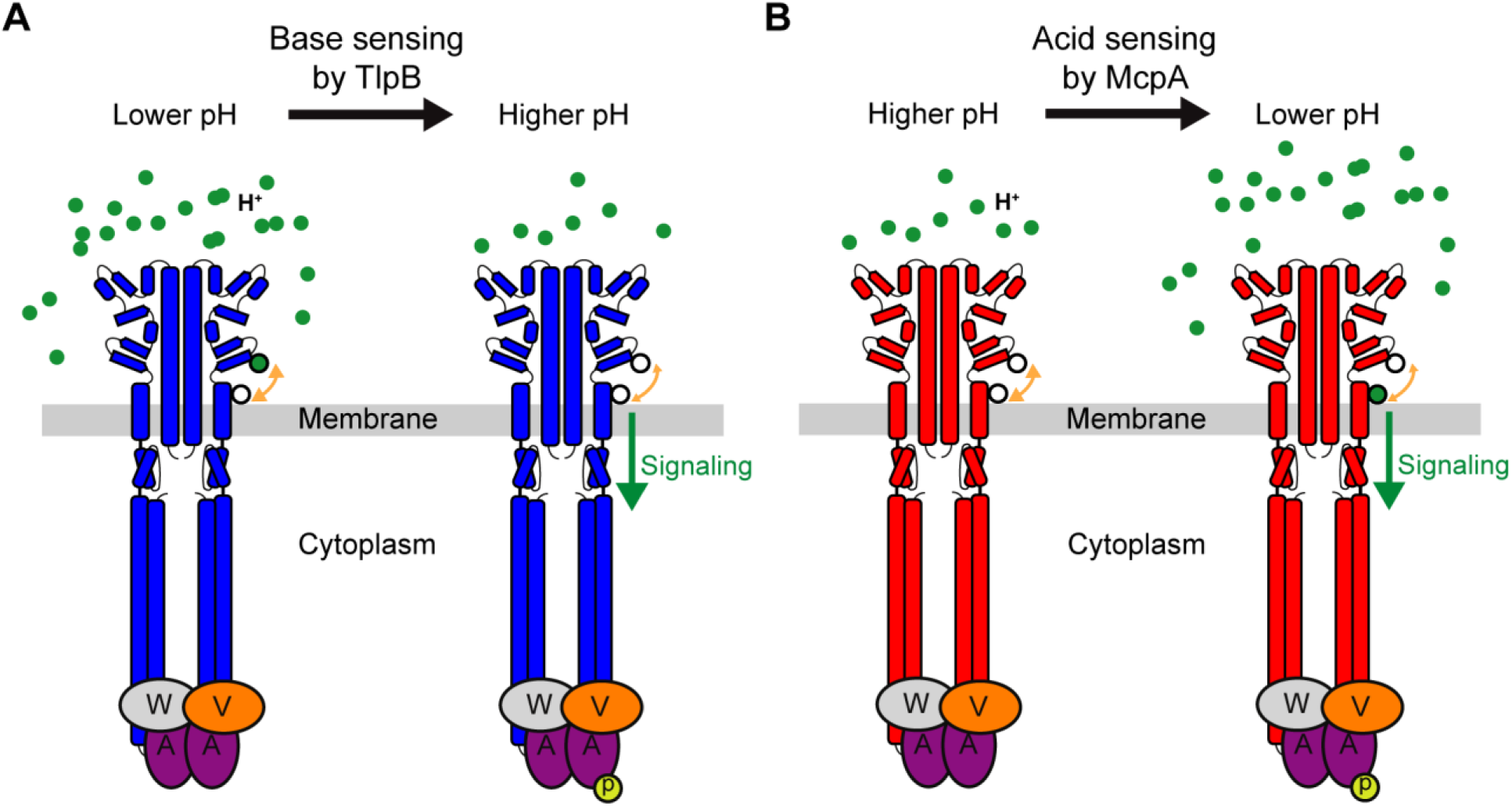
Model for pH sensing mechanism in *B. subtilis.* (**A)** At low pH, two ionizable residues (solid green circle) on TlpB are in their protonated state and form hydrogen bonds with two adjacent residues (white circle). Deprotonation of these residues upon pH increase disrupts the local structure due to decreased hydrogen bonding and induces signaling. (**B**) At high pH, the key histidine residue (lower white circle) within the pH sensing region of McpA is in neutral state and forms hydrogen bonds with adjacent residues (upper white circle). As pH decreases, the histidine residue becomes protonated (solid green circle), leading to loss of hydrogen bonding. This disrupts the local structure and induces signaling.

In the case of McpA, the ionizable residues are His^273^ and Glu^274^. The other two key residues Thr^199^ and Gln^200^ contain polar side-chains that are insensitive to pH changes. As expected, the double mutant H273Q/E274D significantly reduced the response to decrease in pH (**Figure 5D**). Among the two residues, His^273^ seems to play a pivotal role as the point mutation E274Q did not affect the response to pH gradients (1,176.0 ± 124.5 versus 910.4 ± 77.0 cells for wild-type *tlpB*). On possible model is that at high pH (*e.g.* pH=8), His^273^ is in its neutral form. As pH decreases, His^273^ becomes protonated and no longer forms hydrogen bonds with either Thr^199^ and/or Gln^200^. This destabilizes the local structure. Similar to the TlpB case, this conformational change induces a structural transition in signaling module, which in turn promotes phosphorylation of the CheA histidine kinase (**Figure 6B**).

## CONCLUSION

We identified the chemoreceptors governing pH taxis in *B. subtilis*. McpA is the primary acid chemoreceptor while McpB and TlpB are the base chemoreceptors. In addition, TlpA alone functions both as an acid and base chemoreceptor, though its primary role appears to be acid sensing. Using receptor chimeras, we identified four critical amino-acid residues involved in pH sensing. Swapping these residues between McpA and TlpB was able to convert the former into a base sensor and the latter into an acid sensor. Based on our results, we were able to propose a model for pH sensing in *B. subtilis*. Collectively, these results further our understanding of pH taxis and provide a model for pH sensing.

## MATERIALS AND METHODS

### Chemicals and growth media

The following media were used for cell growth: tryptone broth (TB: 1% tryptone and 0.5% NaCl); tryptose blood agar base (TBAB: 1% tryptone, 0.3% beef extract, 0.5% NaCl, and 1.5% agar); and capillary assay minimal medium (CAMM: 50 mM potassium phosphate buffer (pH 7.0), 1.2 mM MgCl_2_, 0.14 mM CaCl_2_, 1 mM (NH_4_)_2_SO_4_, 0.01 mM MnCl_2_, and 4.2 µM ferric citrate). Chemotaxis buffer consists of 10 mM potassium phosphate buffer (pH 7.0), 0.14 mM CaCl_2_, 0.3 mM (NH_4_)_2_SO_2_, 0.1 mM EDTA, 5 mM sodium lactate, and 0.05% (v/v) glycerol.

### Bacterial strains and plasmids

All *B. subtilis* strains were derived from the strain OI1085 (38). All cloning was performed using NEB® 5-alpha Competent *E. coli* (New England Biolabs). Bacterial strains and plasmids used in this work are listed in **Tables 1** and **2**, respectively. All oligonucleotides used in this study are provided in **Table S1**.

**Table 1.**
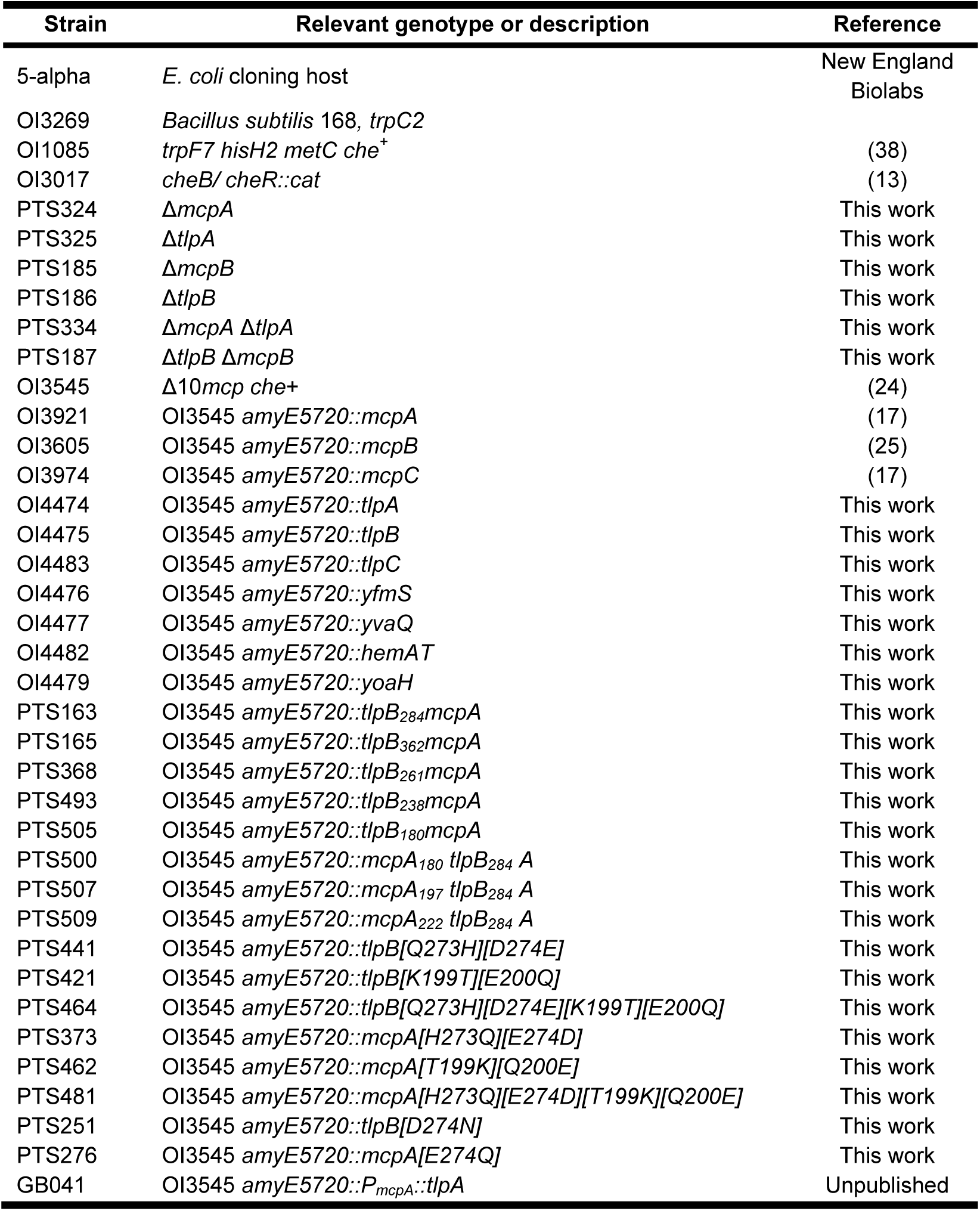
Strains used in this study

**Table 2.** Plasmids used in this study

| Plasmid | Description | Reference |
| --- | --- | --- |
| pJOE8999 | Shuttle vector for Cas9 expression and tracrRNA transcription; Kan<sup>R</sup> | (39) |
| pJSpe | Modified pJOE8999 optimized for Gibson assembly of homology templates; Kan<sup>R</sup> | This work |
| pPT058 | pJSpe::*mcpB* (*mcpB* knockout vector) | This work |
| pPT074 | pJSpe::*tlpB* (*tlpB* knockout vector) | This work |
| pPT116 | pJSpe::*mcpA* (*mcpA* knockout vector) | This work |
| pPT118 | pJSpe::*tlpA* (*tlpA* knockout vector) | This work |
| pPT141 | pJSpe::*mcpA-tlpA* (*mcpA-tlpA* knockout vector) | This work |
| pAIN750 | *B. subtilis amyE* integration vector; Amp<sup>R</sup>, Spc<sup>R</sup> | (17) |
| pAIN750mcpA | pAIN750::*mcpA* | (25) |
| pAIN750mcpB | pAIN750::*mcpB* | (25) |
| pAIN750mcpC | pAIN750::*mcpC* | (17) |
| pAIN750tlpA | pAIN750::*tlpA* | This work |
| pAIN750tlpB | pAIN750::*tlpB* | This work |
| pAIN750tlpC | pAIN750::*tlpC* | This work |
| pAIN750yfmS | pAIN750::*yfmS* | This work |
| pAIN750yvaQ | pAIN750::*yvaQ* | This work |
| pAIN750hemAT | pAIN750::*hemAT* | This work |
| pAIN750yoaH | pAIN750::*yoaH* | This work |
| pPT065 | pAIN750::*tlpB*<sub>362</sub>*mcpA* | This work |
| pPT063 | pAIN750::*tlpB*<sub>284</sub>*mcpA* | This work |
| pPT143 | pAIN750::*tlpB*<sub>261</sub>*mcpA* | This work |
| pPT224 | pAIN750::*tlpB*<sub>238</sub>*mcpA* | This work |
| pPT234 | pAIN750::*tlpB*<sub>180</sub>*mcpA* | This work |
| pPT233 | pAIN750::*mcpA*<sub>180</sub> *tlpB*<sub>284</sub> *A* | This work |
| pPT236 | pAIN750::*mcpA*<sub>197</sub> *tlpB*<sub>284</sub> *A* | This work |
| pPT237 | pAIN750::*mcpA*<sub>222</sub> *tlpB*<sub>284</sub> *A* | This work |
| pPT129 | pAIN750::*tlpB*[Q273H][D274E] | This work |
| pPT162 | pAIN750::*tlpB*[K199T][E200Q] | This work |
| pPT202 | pAIN750::*tlpB*[Q273H][D274E][K199T][E200Q] | This work |
| pPT196 | pAIN750::*mcpA*[H273Q][E274D] | This work |
| pPT163 | pAIN750::*mcpA*[T199K][Q200E] | This work |
| pPT222 | pAIN750::*mcpA*[H273Q][E274D][T199K][Q200E] | This work |
| pPT101 | pAIN750::*tlpB*[D274N] | This work |
| pPT107 | pAIN750::*mcpA*[E274Q] | This work |
| pGB045 | pAIN750::P<sub>mcpA</sub>::*tlpA* | Unpublished |

Gene deletions were constructed using plasmids derived from pJSpe. pJSpe was derived from pJOE8999, which provides a CRISPR/Cas9-based, marker-free genome editing system for *B. subtilis* (39). We found that the *SfiI* restriction sites on the original pJOE8999 were inefficient for subcloning homology templates. In addition, a 13-bp long DNA scar remained on the chromosome after the targeted DNA fragment was deleted using pJOE8999-derived vectors. Therefore, we created pJSpe for more efficient assembly of homology templates based on Gibson assembly and scar-less deletion of DNA fragments. Briefly, a 50-bp annealed complementary DNA oligonucleotides containing a *SpeI* restriction site and optimized for Gibson assembly was inserted between the *SfiI* restriction sites on pJOE8999 to yield pJSpe. For construction of the knockout plasmids, a 20-bp sgRNA target sequence for the targeted gene was designed using CRISPy-web online tool (40). Annealed complementary oligonucleotides were then subcloned into *BsaI* restriction sites on pJSpe as described in (39). Two PCR fragments (∼600-800 bp) flanking the targeted gene using overlapping primers were subcloned into *SpeI-*linearized vector using Gibson assembly (41). The resultant vector was then linearized using *XhoI* and ligated with T4 DNA ligase to create a long DNA concatemer. The concatemer was then transformed into *B. subtilis* strain using the two-step Spizizen method (42). Single colonies were isolated and twice streaked on fresh plates. Plasmid curing and genotype verification were performed as previously described (39).

Strains expressing a single wild-type chemoreceptor were constructed by integrating the respective chemoreceptor expression cassette into the *amyE* locus. The region containing the promoter, gene, and terminator was PCR amplified from genomic DNA isolated from *B. subtilis* 168. The PCR fragment was then cloned into the plasmid pAIN750 using the *EcoRI* and *BamHI* restriction sites. The plasmid was then transformed in OI3545, which lacks all ten chemoreceptors, as described above.

Chemoreceptor chimeras were constructed using Gibson assembly (41). Briefly, two opposing primers were designed to prime DNA synthesis outwards from the fusion point of the chimeric gene using PCR with pAIN750*mcpA* as the DNA template. Then, a second pair of primers with overlapping regions were designed to PCR-amplify the desired fragment of *tlpB* gene from pAIN750*tlpB* plasmid. Following purification the PCR products by gel extraction, the DNA fragments were assembled (41).

Point mutations were introduced into chemoreceptor genes using the inverse PCR method with pAIN750*mcpA* and pAIN750*tlpB* as DNA templates. Briefly, plasmid was PCR amplified using two opposing primers containing the desired mutations. Following purification by gel extraction, the DNA fragment was phosphorylated with T4 polynucleotide kinase and then blunt-end ligated using T4 DNA ligase. Ligation product was heat-inactivated and transformed into *E. coli*. The plasmid was then isolated from*E. coli* and transformed into *B. subtilis* OI3545 as described above.

### Capillary assay for chemotaxis

The capillary assay was performed as described previously (1, 26). Briefly, cells were grown for 16 hours at 30 °C on TBAB plates. The cells were then scraped from the plates and resuspended to *A*_600_ = 0.03 in 5-mL CAMM supplemented with 50 μg/mL histidine, methionine, tryptophan and 20 mM sorbitol, and 2% TB. The cultures were grown to *A*_600_ = 0.4-0.45 at 37 °C and 250 rpm shaking. At this point, 50 μl of GL solution (5% (v/v) glycerol, 0.5 M sodium lactate) was added to culture and the cells were incubated for another 15 minutes (at 37 °C, 250 rpm shaking). Cells were then washed twice with chemotaxis buffer (pH 7.0) and incubated for additional 25 minutes (at 37 °C, 250 rpm shaking) to assure that the cells were motile. Cells were diluted to *A*_600_ = 0.001 in chemotaxis buffer at desired pH values for pH taxis experiments and in chemotaxis buffer (pH 7.0) for casamino-acid control experiments. For indole control experiments, cells were diluted to *A*_600_ = 0.005 in chemotaxis buffer containing indole (50 μM, pH 7.0). Prior to assay, cells were briefly incubated in chemotaxis buffer at room temperature (shaking slowly) and then aliquoted into 0.3-mL ponds on a slide warmer at 37 °C and closed-end capillary tubes filled with chemotaxis buffer or casamino acid (3.16 x 10^−5^ g/mL) solutions prepared with chemotaxis buffer (pH 7.0) were inserted. After a fixed time (30 minutes for casamino acids and 1 hour for pH and indole), cells that migrated into the capillaries were harvested and transferred to 3 mL of top agar (1% tryptone, 0.8% NaCl, 0.8% agar, 0.5 mM EDTA) and plated onto TB (1.5% agar) plates. These plates were incubated at 37 °C for 16 hours and colonies were counted. Experiments were performed in triplicate each day and then repeated on three different days.

### Structural analysis

Domain predictions were performed using he phmmer search engine on the HMMER web-server using the UniProt reference proteomes database with default sequence E-values thresholds (43). Boundaries of both transmembrane alpha helices were first predicted using TMHMM web-server v.2.0 (44) and then manually adjusted using information from propensity analysis of amino acid distributions around lipid/water interfaces (45). Pairwise amino-acid sequence alignments between McpA and TlpB for chimeric-receptor analysis were performed using EMBOSS Water (46) and multiple sequence alignment between McpA, McpB, TlpA, and TlpB for mutational analysis were carried out in T-Coffee (47). Structures for the McpA and TlpB sensing domains (residues 38-278) were predicted using the I-TASSER web-server (48). The C-scores were 1.15 and 1.13 for McpA and TlpB, respectively. Both models are structurally close to the ligand-binding domain of the PctA chemoreceptor from *Pseudommonas aeruginosa PAO1* (PDB: 5LTX) with TM-score of 0.955 for both McpA and TlpB. Visualization of all structures was accomplished using the VMD software package (v-1.9.3) (49).

## ACKNOWLEDGEMENTS

This work was supported by the University of Illinois through the Robert W. Schaefer Faculty Scholar fund and National Institutes of Health Grant GM054365.

